# Spatiotemporal regulation of Pseudorabies Virus Thymidine kinase UL23 by post-translational modification

**DOI:** 10.1101/2024.03.06.583799

**Authors:** Chuang Li, Rui Cao, Rui Zhang, Jun Tang

## Abstract

Post-translational modification plays a significant role in the interaction between viruses and their hosts. The pseudorabies virus (PRV) is a highly contagious herpesvirus that affects the central nervous system and respiratory tract of swine. However, the role of post-translational modifications, including Ubiquitination and SUMOylation, in host and PRV interplays is poorly understood. Here we examined the SUMO modification of PRV proteins and revealed that the PRV thymidine kinase UL23 can undergo SUMO modification. Bioinformatic analysis suggested four potential modification sites for UL23. Site-directed mutagenesis indicates that SUMO modification occurs at sites K242 and K267 as the wild type localizes in the nucleus while the mutants localize in the cytoplasm. Subsequently, polyclonal antibodies against mouse derived UL23 were employed to reveal that wild type UL23 was mainly located in the nucleus during PRV infection. Co-expression of UL23 with SUMO deconjugating enzymes showed that SENP1/2 inhibited the nuclear import of UL23. Whereas SUMO modification significantly impacted the localization of UL23, it did not detectably affect its stability.

## Introduction

The Pseudorabies virus (PRV) is the causative agent of pseudorabies in swine^1^, belonging to the alpha-herpesvirus family, can infect a variety of domestic and wild animals, causing pseudorabies with pigs serving as the natural host and reservoir of the virus^2^. The alpha-herpesvirus family also includes viruses that can cause human diseases such as Herpes simplex virus (HSV)-1, HSV-2, and Varicella-zoster virus (VZV)^3^. To date, there have been no confirmed reports of pseudorabies in humans; however, corresponding antibodies can be detected in human serum. Pseudorabies outbreaks and epidemics in major swine-raising countries worldwide have caused significant economic losses to the swine farming industry^4^. Many countries and regions have invested substantial financial and human resources into research on the disease, hoping to find definitive measures for its prevention, control, and eradication^5^. In some developed countries, effective outcomes have been achieved, with the eradication or control of the pseudorabies virus^6^.

Not only agricultural experts but also neurobiologists and virologists have shown a great interest in the study of the pseudorabies virus^7^. PRV has been used as a model virus for studying the molecular biology of the human herpesvirus HSV-1^8^. The UL23 protein is a thymidine kinase^9^ that catalyzes the synthesis of viral DNA, participates in virus replication, and the spread of the virus in the central nervous system^10^. It has been reported that PRV lacking thymidine kinase UL23 exhibits significantly reduced virulence in infected swine^11^. Therefore, *Ul23*-deficient PRV does not affect the immune response and can be used for vaccine preparation^12^. Vaccines derived from UL23-deficient PRV, when administered to swine^13^, do not cause clinical symptoms and can protect animals from PRV infection^14^.

Post-translational modification by the Ubiquitination^15^ and Small Ubiquitin-like Modifier (SUMO) protein family is an important regulatory mechanism for many cellular proteins^16^. Different from Ubiquitin, SUMO molecules have three isoforms: SUMO-2 and SUMO-3 share a high degree of homology, at 95%^17^, whereas SUMO-1 only has 47% homology with SUMO-2/3^18^. Due to their ability to undergo SUMO modification themselves, SUMO-2 and SUMO-3 can form chains, suggesting potential functional differences between SUMO-1 and SUMO-2/3 modifications^19^.

Obligate parasitic viruses must interact with the entire replication cycle of host cells^20^, leading to numerous instances where viral proteins engage in interactions with host protein modification systems^21^. As part of their intracellular lifecycle, viruses employ a variety of proteins to interact with the host, maintaining their advantage while disrupting cellular pathways, yet host cells possess mechanisms to evade antiviral defenses^22^. The Ebola Zaire virus, for example, can suppress the production of type I interferons in the host through SUMO modification^23^. In the case of Hepatitis C Virus (HCV), SUMO modification of IE180 can increase the production of IE2, thereby promoting viral replication, although SUMO modification of IE2 does not affect the virus’s replication in vitro^24^. For HCV IE2 p86, SUMO modification facilitates virus infection. SUMO modification of HCV’s IE1 can alter its cellular localization^25^.

The activity and subcellular localization of viral proteins can be altered by SUMO modification^26^. Nevertheless, role of SUMOylation on PRV proteins have not been reported yet. Thus, we screened PRV proteins for modification by SUMO molecules to further explore the impact of SUMOylation on PRV viral protein functions. PRV has not been eradicated and continues to plague the domestic livestock industry^27^. The vaccine widely used worldwide lacks the thymidine kinase, which is encoded by the *Ul23* gene^12^. This suggests that research on the UL23 SUMOylation could provide a theoretical basis for the development of new vaccines^10^.

## Results

### 1. PRV protein UL23 is SUMOylated *in vitro*

Preliminary experiments have successfully constructed several Pseudorabies virus proteins^28^. Based on this achievement, five proteins US1, US3, UL13, UL23, and UL48 with successful ectopic expression were selected for SUMO modification analysis. These proteins were co-transfected with SUMO1, the SUMO E2 conjugating enzyme UBC9. PML4, which is known to undergo SUMO modification, serves as a positive control.

After purifying FLAG-UL23 protein with M2 beads, HA-SUMO1 was identified using an HA antibody in **Figure 1A**, showing that UL23, along with PML4, can undergo SUMO modification. In **Figure 1B**, HA-SUMO1 was purified with HA beads, and the viral proteins tagged with FLAG were identified using a FLAG antibody, revealing that only UL23 and PML4 can be modified by SUMO1. There are five known SUMO isoforms, with the first three being well-known and the fifth discovered in early 2016; different isoforms of SUMO have distinct functions^29^. To examine the SUMO modification of UL23 protein by these three proteins, FLAG-UL23 was co-transfected with HA-SUMO1/2/3. **Figure 1C** shows that UL23 can be modified by all three isoforms of SUMO proteins. In **Figure 1D**, FLAG-UL23 was transfected into 293T cells alone to detect the modification of UL23 protein by endogenous SUMO1, identified using an antibody against SUMO1. It was observed that UL23 protein can be modified by endogenous SUMO1.

**Figure 1:**
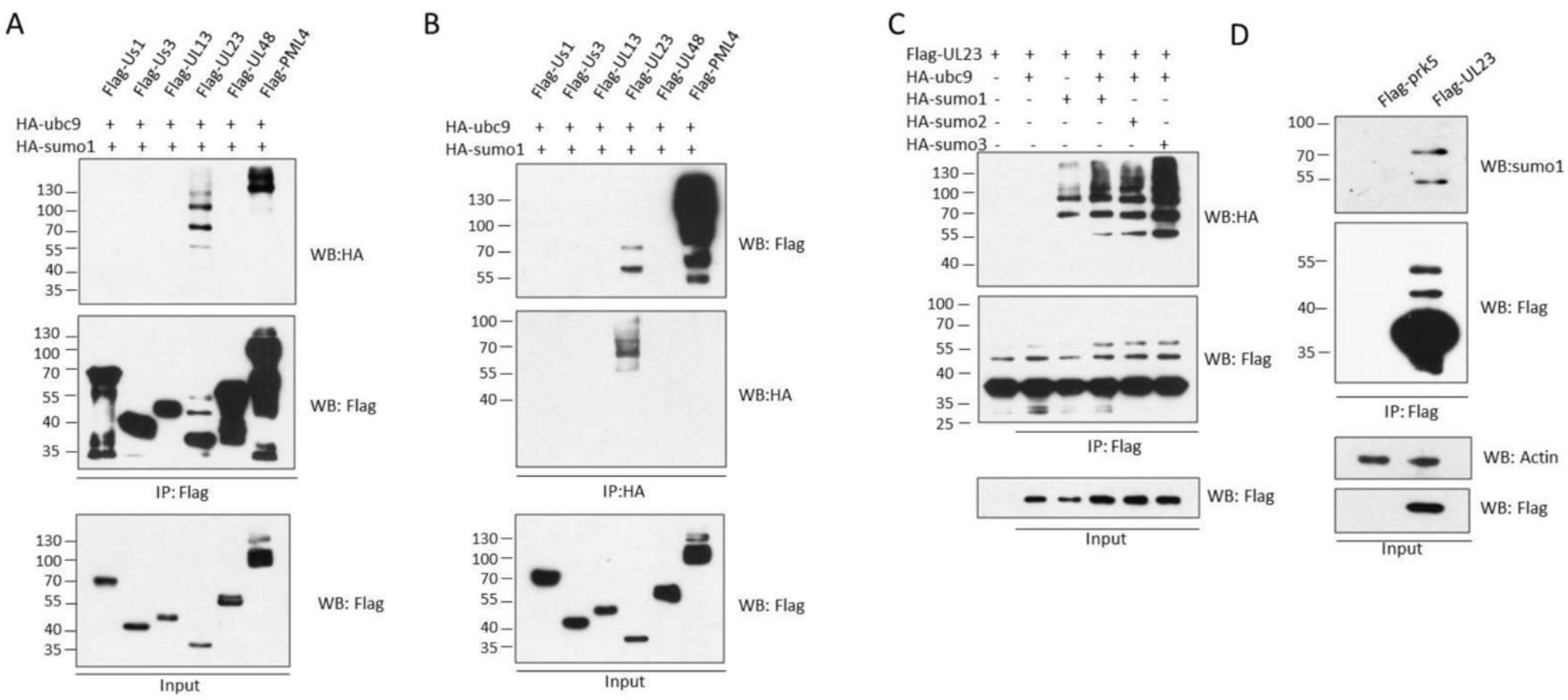
SUMOylation of PRV protein UL23.

### 2. Identification of SUMOylation sites on PRV UL23

Protein SUMOylation primarily occurs on Lysine residues, and mutating the Lysine to Arginine abolishes SUMO modification. Through analysis of the UL23 amino acid sequences, we hypothesized four potential SUMO modification sites: K16, K57, K242, and K267, as shown in **Figure 2A**. After generating mutants of UL23 with Lysine individually mutated to Arginine using site-directed mutagenesis^30^, we examined the SUMO modification of these mutants. **Figure 2B** reveals that the SUMO modification levels of K16R and K57R mutants are similar to the wild type, whereas K242R and K267R mutants show significantly lower SUMO modification levels compared to WT. This suggests that SUMO modification of UL23 likely occurs at K242 and K267. Subsequent construction of mutants containing both K242 and K267 mutations, as well as a mutant with all four Lysines mutated, was carried out. **Figure 2C** indicates that these mutants exhibit significantly reduced SUMOylation compared to WT, leading to the conclusion that SUMO modification of the pseudorabies virus UL23 protein occurs at K242 and K267.

**Figure 2:**
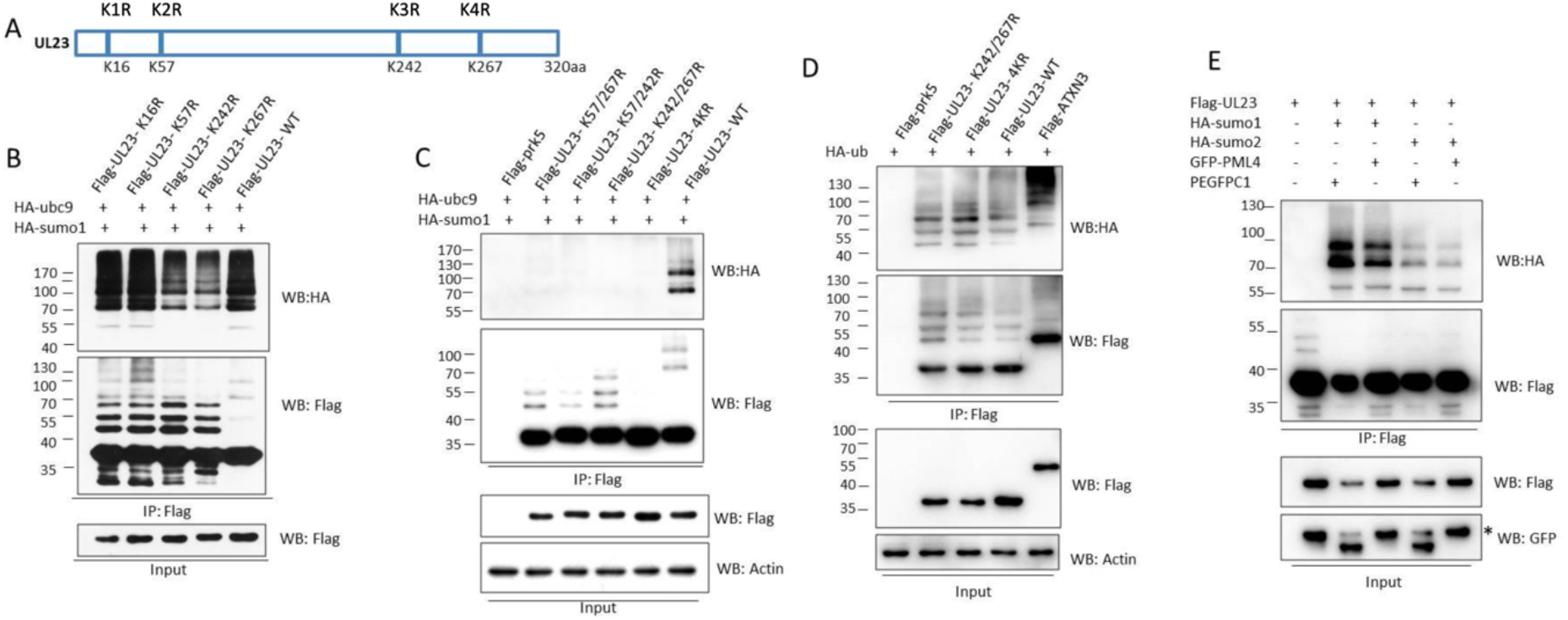
Identification of SUMOylation sites on PRV UL23.

Further inspection of the immunoprecipitation (IP) results reveals upward shifts in some UL23 protein bands following K to R mutations, a pattern similar to that seen with ubiquitinated proteins^31^. Consequently, we co-transfected FLAG-UL23 and HA-ubiquitin to assess the ubiquitination status of UL23, using ATXN3, which is known to undergo ubiquitination, as a positive control. **Figure 2D** shows that both the WT and mutant forms of UL23 can undergo ubiquitination. Since PML, a major component of PML nuclear bodies, plays a crucial role in regulating intracellular SUMO modifications of many proteins, we aimed to investigate whether PML4 could influence the SUMOylation of UL23. Thus, FLAG-UL23, HA-SUMO1/2, and GFP-PML4 were co-transfected to examine the SUMOylation status of UL23. **Figure 2E** demonstrates that PML4 does not affect the degree of SUMO1/2 modification of UL23.

### 3. Sub-cellular localization of PRV UL23

For viral proteins to function effectively, correct cellular localization is essential, with subcellular localization generally categorized into nucleus, nuclear membrane, cytoplasm, and cell membrane. Our initial goal was to determine the cellular localization of the wild-type UL23 protein during PRV infection of cells. Using PRV to infect PK15 cells and, after 12 hours, employing homemade UL23 antibodies to identify PRV’s UL23 protein, **Figure 3A** reveals that the majority of the wild-type UL23 protein is localized in the nucleus, with a minor amount in the cytoplasm. US3, which localizes to the nucleus, served as a positive control for this experiment^32^. We transfected various mutants of FLAG-UL23-prk5 into Hela cells and observed their subcellular localization through immunofluorescence, as illustrated: **Figure 3B** shows that the WT UL23 protein localizes to the nucleus, the K16 lysine to arginine mutation (K1R) still localizes within the nucleus, forming punctate structures, and K57 lysine mutation does not affect the localization of UL23 in the nucleus. However, mutations at K242 or K267 (K3R/K4R) result in a portion of UL23 protein localized in the cytoplasm, also forming punctate aggregates within the cytoplasm. This preliminary conclusion suggests that lysine mutations at K242 and K267 affect the subcellular localization of UL23.

**Figure 3:**
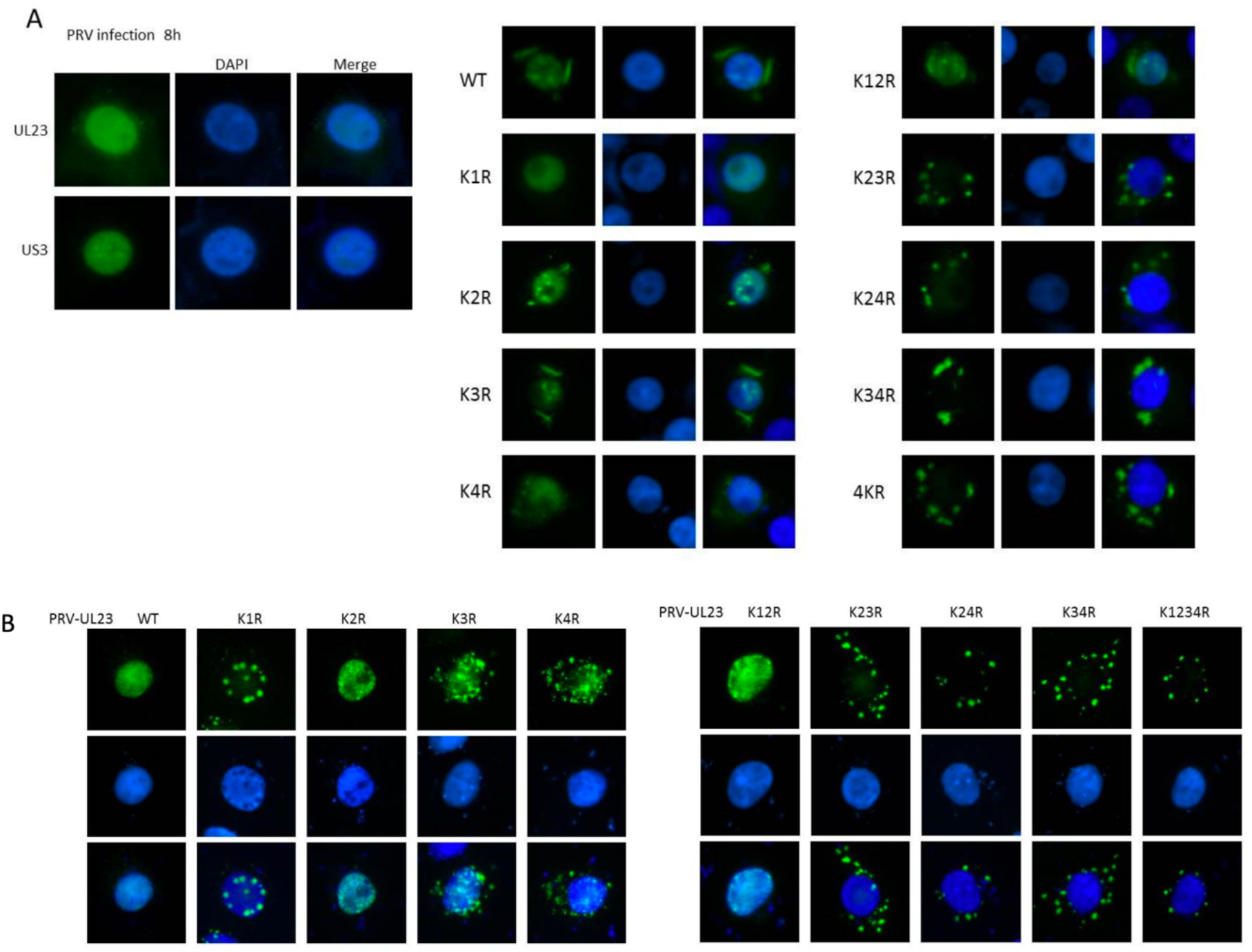
Sub-cellular localization of PRV UL23.

To identify the amino acids affecting UL23 localization, we constructed double and quadruple mutant UL23 variants on the basis of previously mutated Lysines. **Figure 3C** indicates that the K16/57R mutation (K12R) does not affect the localization of UL23 in the nucleus, whereas the K57/242R, K57/267R, K242/267R, and K16/57/242/267R mutants, which include mutations at the third and fourth lysine positions, primarily localize in the cytoplasm and are distributed in a punctate manner. This further confirms the sites affecting UL23 localization.

### 4. Impacts of SENPs on the localization of UL23

To further verify our experimental results, PK15 cells were infected with PRV, and samples were collected at 0 hr, 12 hr, and 24 hr post-infection. Subsequently, the obtained samples underwent nuclei-cytoplasmic separation to measure the abundancy of UL23 protein in both the cytoplasm and the nucleus. As shown in **Figure 4A**, PRV UL23 protein is predominantly located in the nuclear compartment. Tubulin serves as a cytoplasmic marker while H3 acts as a nuclear marker^33^. Due to the absence of PRV viruses with UL23 protein SUMOylation site mutations, we were forced to investigate the subcellular localization of exogenous UL23 protein wild-type and mutants. FLAG-UL23-WT, FLAG-UL23-K34R, and FLAG-UL23-4KR were transfected into Hela cells, followed by sample collection and nuclei-cytoplasmic separation. It was observed from **Figure 4B** that the wild-type WT is mainly located in the nuclear compartment, while the SUMOylation mutants K3/4R or 4KR are predominantly found in the cytoplasmic compartment. Laminb, locates and functions in the nucleus, can also serve as a nuclear marker^34^. From these conclusions, it is apparent that the wild-type UL23 protein is primarily localized in the nucleus, while the SUMOylation mutated UL23 proteins are mainly localized in the cytoplasm.

**Figure 4:**
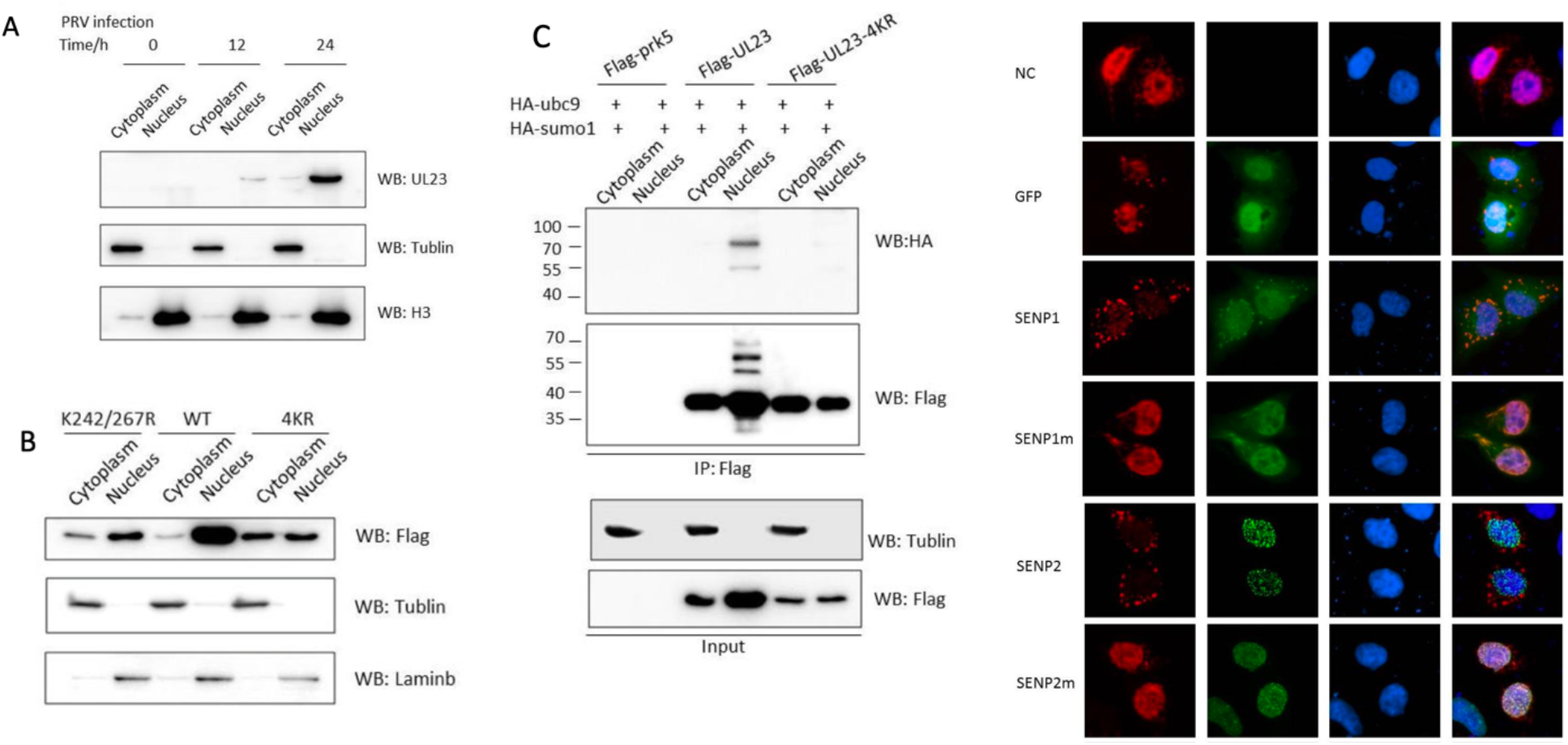
Impacts of SENPs on the localization of UL23.

To further validate the impact of SUMOylation on the localization of UL23, the SUMOylation status of UL23-WT and 4KR in separated nuclear and cytoplasmic fractions was examined. As shown in **Figure 4C**, only the UL23 protein located in the nucleus undergoes SUMOylation, indicating that only SUMOylated UL23 protein can be localized in the nucleus.

SUMOylation is a dynamic and reversible process, regulated in two main aspects: covalent attachment mediated by SUMO ligases and de-SUMOylation regulated by a class of de-SUMOylating enzymes, sentrin-specific proteases (SENPs). If the SUMOylation mutants can affect the subcellular localization of UL23 protein, then the SENP enzymes that remove SUMOylation should have a similar effect. To test this hypothesis, SENPs and UL23 were co-transfected into Hela cells to observe the effect of SENPs on the subcellular localization of UL23 protein. As shown in **Figure 4D**, transfection of UL23 alone results in nuclear localization; GFP does not affect the localization of UL23, while SENP1 and SENP2 can hinder UL23 nuclear entry. Mutation of the active sites of SENP1 and SENP2’s de-SUMOylating enzyme activity no longer affects UL23 nuclear entry. In addition, SENP3 does not affect the localization of UL23.

## Discussion

The occurrence of SUMOylation in viral proteins is not surprising, as numerous studies have reported the crucial role of SUMOylation in the interactions between viruses and their hosts^26^. Currently, the Pseudorabies virus still poses a threat to the livestock industry worldwide, causing significant damage to agriculture and the economy^5^. Previous research primarily focused on viruses infecting humans, with relatively less attention paid to those infecting animals^35^, except for notable studies on avian influenza and porcine reproductive and respiratory syndrome virus^36(p402)^. In this study, we discovered that the UL23 protein of PRV can undergo SUMOylation and further explored the effects of SUMOylation on the localization of UL23 protein.

Through immunoprecipitation, we validated the interaction between SUMO proteins and the UL23 protein from two different perspectives, confirming that UL23 can be modified by SUMOylation. There are three different isoforms of SUMO proteins: SUMO-1/2/3, and it was found that UL23 can be modified by all three isoforms^37^. Whether the functions of UL23 modified by different SUMO protein isoforms are consistent remains to be investigated. After analyzing the amino acids of UL23, we predicted four potential SUMOylation sites^38^. Mutating these sites individually revealed that the lysine mutations at K242 and K267 resulted in decreased SUMOylation levels. Further construction of a UL23 protein with multiple mutations confirmed that SUMOylation of UL23 indeed occurs at K242 and K267.

SUMOylation can regulate the localization of proteins, and in this context, we aimed to study whether SUMOylation could affect the localization of the UL23 protein. Using a homemade UL23 antibody to recognize UL23 protein expressed in PRV-infected PK15 cells, we found that the wild-type UL23 protein is primarily located in the nucleus^39^. In our experiments, mutating lysine (K) to arginine (R) affected the site’s SUMOylation but did not alter its nuclear localization signal^40^. After comparing the subcellular localization of the wild-type UL23 protein with that of the SUMOylation mutant, we observed that the wild-type mainly localized in the nucleus, while the SUMOylation mutant was primarily located in the cytoplasm. This suggests that SUMOylation indeed significantly influences the localization of UL23.

To further confirm that the cytoplasmic localization of UL23 was not due to changes in the nuclear localization signal, we co-transfected the de-SUMOylating enzyme SENP with UL23 into Hela cells to observe the effect of SENP on the subcellular localization of UL23 protein. It was found that transfecting UL23 alone results in nuclear localization; GFP does not affect the localization of UL23, while enzymatic active SENP1 and SENP2 can inhibit UL23 nuclear entry.

In addition to regulating the subcellular localization of proteins, SUMOylation can also modulate protein stability and activity^41^. In this study, we aimed to investigate whether SUMOylation could affect the stability of the UL23 protein. After treating with cycloheximide (CHX) and collecting samples at different time points to measure UL23 protein levels, we found that there was no significant difference in the half-life between the wild type and the SUMOylation mutant. To further determine the impact of SUMOylation on the stability of UL23, we inhibited proteasomal degradation by pre-treating with MG132 before collecting samples, to prevent the degradation of UL23 protein. The results showed no significant difference between the negative control group and the MG132-treated group, indicating that SUMOylation does not affect the stability of UL23 protein.

In our comprehensive study, we investigated the effects of SUMOylation on the PRV UL23 protein, focusing on its localization^42^. Our efforts discovered that UL23 undergoes SUMOylation and identified specific lysine residues essential for this modification. Our findings revealed that SUMOylation significantly influences the subcellular localization of UL23, with the wild-type protein predominantly located in the nucleus and SUMOylation mutants primarily in the cytoplasm^43^. This suggests that SUMOylation plays a critical role in the nuclear localization of UL23. However, our experiments showed that SUMOylation does not affect the stability of the UL23 protein, as evidenced by similar degradation rates between the wild type and SUMOylation mutants, even when the proteasomal degradation pathway was inhibited using MG132^44^. This research highlights the multifaceted role of SUMOylation in regulating viral protein function, particularly in terms of localization, without impacting protein stability.

